# Grass fly evolution unlocked: phylogenomics and classification of worldwide Chloropidae (Diptera)

**DOI:** 10.1101/2025.09.30.679457

**Authors:** Paula Raile Riccardi, Dalton de Souza Amorim, Keith Bayless, Joshua V. Peñalba

## Abstract

Chloropidae, commonly known as grass flies, is a hyperdiverse family of true flies thriving in terrestrial ecosystems around the world with importance to conservation and economy. However, true grass fly diversity and evolutionary affinities are still largely unknown leading to an obscure classification at varied taxonomic levels. The massive lack of data on grass fly evolution hampers studies ranging from species discovery to diversification and provision of ecological services. To overcome this issue, we provide the first comprehensive phylogeny of worldwide Chloropidae using whole-genome shotgun sequencing to retrieve mitochondrial genes, ultraconserved elements and single-copy orthologs. With this, we generated genomic resources from all subfamilies, 95% of the tribes and 48% of the chloropid genera from all biogeographic regions which provides an invaluable resource for biodiversity studies worldwide. Overall, our results provide novel genomic data of over 100 fly species. We implemented dramatic changes in grass fly classification system that redefine one subfamily, propose three new tribes, reassign the tribal position of 59 genera, and synonymize four suprageneric taxa. In addition, we reveal a novel synapomorphy for the family, elucidate grass fly sister-group, and assess the utility of the mitochondrial genome alone to recover evolutionary relationships of a recent lineages of flies. This study is fundamental to mitigate the neglect associated with grass flies through an unprecedented source of genomic data, ultimately providing a novel framework for understanding the rapid evolution of the last and most species-rich radiation of flies.

## 1. INTRODUCTION

Chloropidae, commonly known as grass flies, is a hyperdiverse family of flies with over 3,000 described species in about 200 genera thriving in virtually all terrestrial ecosystems (Riccardi & Hartop 2024). They are particularly abundant in open habitats (Riccardi & Hartop 2024; Andersson & Nartshuk 2013). From agricultural pests to gall makers, disease vectors, parasites, and even pollinators, chloropids play a diverse and crucial role in ecosystems around the world.

This lineage, as well as the other schizophoran flies, was originated after the K–T extinction event, being part of the most species-rich radiation of animal life that comprise a third of fly diversity (Bayless et al 2021). Bayless et al. (2021) clarified the evolutionary relationships among many schizophoran families, but the position of chloropids were still left unresolved.

These large gaps in knowledge about grass fly diversity and evolutionary history lead to an obscure classification system at varied taxonomic levels (Riccardi & Amorim 2020). Their understudied morphology and scarce genetic information have hindered our understanding of their species richness, distribution, and evolutionary history. This fact further limits studies ranging from species discovery to diversification and provision of ecological services.

Historically, the relationships among grass flies proposed mainly by Andersson (1977) and Nartshuk (1984, 2012) do not have an explicit set of arguments nor a cladistic framework as they mostly disregard the endemic tropical fauna. Moreover, Liu et al. (2024) tried to infer phylogenetic relationships based on mitochondrial genomes including only 12 genera and 14 grass fly species. The later study failed to resolve the relationships between the subfamilies and provided taxonomic sampling too limited for addressing the problematic grass fly classification system.

Chloropids are currently classified into four subfamilies based exclusively on morphology. The subfamily Siphonellopsinae is composed of four genera and is considered sister to the remaining chloropids mainly due to the pilosity pattern on the head and scutum (Andersson 1977; Nartshuk 1984). The subfamily Chloropinae has 76 genera, with several taxa reported feeding on monocots, and it is the only subfamiliar rank with a formal phylogeny (Riccardi & Amorim 2020). The Oscinellinae is the most diverse subfamily in terms of feeding strategies and includes 110 genera (Nartshuk 2012; Riccardi et al. 2018; Mlynarek & Wheeler 2018). Subfamily rank has been recently given to the Rhodesiellinae, which comprises 14 genera (Nartshuk 2012) that were previously placed in the Oscinellinae (Andersson, 1977). While Siphonellopsinae and Chloropinae are well-established ranks, the limits between Oscinellinae and Rhodesiellinae are still dubious (Mynarek & Wheeler 2018; Bazyar 2019). Also, the relationships among all subfamilies are uncertain. Moreover, 57 chloropid genera do not belong to any more inclusive classification rank (eg. tribe), and the limits of several speciose genera are dubious, such as *Chlorops* Meigen, *Tricimba* Lioy (Riccardi & Amorim 2020, Riccardi & Hartop 2024). The establishment of a robust classification system would also facilitate the identification process at varied taxonomic levels.

Recent advances in DNA sequencing technology have dramatically increased the information available for systematic studies. Genomic data is particularly useful for disentangling challenging evolutionary relationships of large radiations of organisms, that can be further complicated by morphological plasticity such as that found in grass flies (Riccardi & Amorim 2020; Bazyar 2019). As the cost of sequencing continue to plummet, it is becoming more practical to sequence genomes at low coverage and extract the desired loci in silico rather than perform genome reduction methods prior to sequencing. This advancement is particularly beneficial for rapidly retrieving genetic markers from fresh and historical samples of non-model and minute organisms (Zhang et al. 2019). Moreover, ultraconserved elements (UCEs), benchmarking universal single-copy orthologs (BUSCOs), and mitochondrial DNA are of known utility and effective to infer phylogenetic relationships for other insect lineages (Rhoden &Wahlberg 2020; Zhang et al. 2019). The largely conserved nature of these markers also lends itself to resolving deeper relationships as well as provides a set of loci that can be used across studies. Although UCEs and BUSCOs have been largely used for other insect orders, there are only three publications leveraging UCEs to recover the evolutionary history of robber flies, dance flies and flesh flies (Cohen et al. 2021; Rhoden &Wahlberg 2020; Buenaventura et al. 2020).

Phylogenetic approaches using molecular data within grass flies are so far restricted to few taxa and mitochondrial genes (Triselyova et al. 2014; Liu et al. 2024), which fail to provide enough information to resolve the evolutionary relationships of this group. Also, a recent analysis of schizophoran flies using transcriptomics (Bayless et al. 2021) challenged the traditional assumption of Chloropidae and Milichiidae as sister-groups within Carnoidea (Buck 2006). However, genomic data have never been extensively used to clarify the affinities within Carnoidea.

To overcome the dubious grass fly classification system and obscure evolutionary affinities within Schizophora, we provide the first comprehensive phylogeny of worldwide Chloropidae using genome scale data of fresh and historical specimens. We use whole-genome shotgun sequencing to leverage protein-coding mitochondrial genes, UCEs and BUSCOs for phylogenetic analyses. With this, we generated genomic resources from all subfamilies, 95% of the tribes and 48% of the chloropid genera from all biogeographic regions which provides a baseline for biodiversity studies worldwide. Overall, our outputs provide novel genomic data for over 100 fly species. We established a robust classification system at subfamily and tribe levels and revealed a novel synapomorphy for the family. Moreover, we elucidated the sister-group of grass flies and assessed the utility of the mitochondrial genome alone to recover evolutionary relationships within a recent lineage of flies. This study is fundamental to mitigate the neglect associated with grass flies through an unprecedented resource of genomic data, ultimately providing a novel framework for understanding the obscure evolution and biodiversity associated to the last radiation of flies.

## 2. MATERIAL & METHODS

### 2.1. Taxonomic sampling

The datasets comprise of 117 individuals including 105 ingroups that cover the taxonomic breadth of Chloropidae (Figure 1). The ingroup taxa represents 95 genera and the only tribe not sampled is Pseudothaumatomyini. We included 18 genera without any tribal classification, considered here as *incertae sedis*. The identification of the specimens was confirmed by PRR. Subfamily, tribe and genus ranks were retrieved from Riccardi & Amorim (2020), Nartshuk (2012), and Mlynarek & Wheeler (2018). The validity of the *incertae sedis* genera was checked in von Tschirnhaus & Groll (2024).

**Figure 1:**
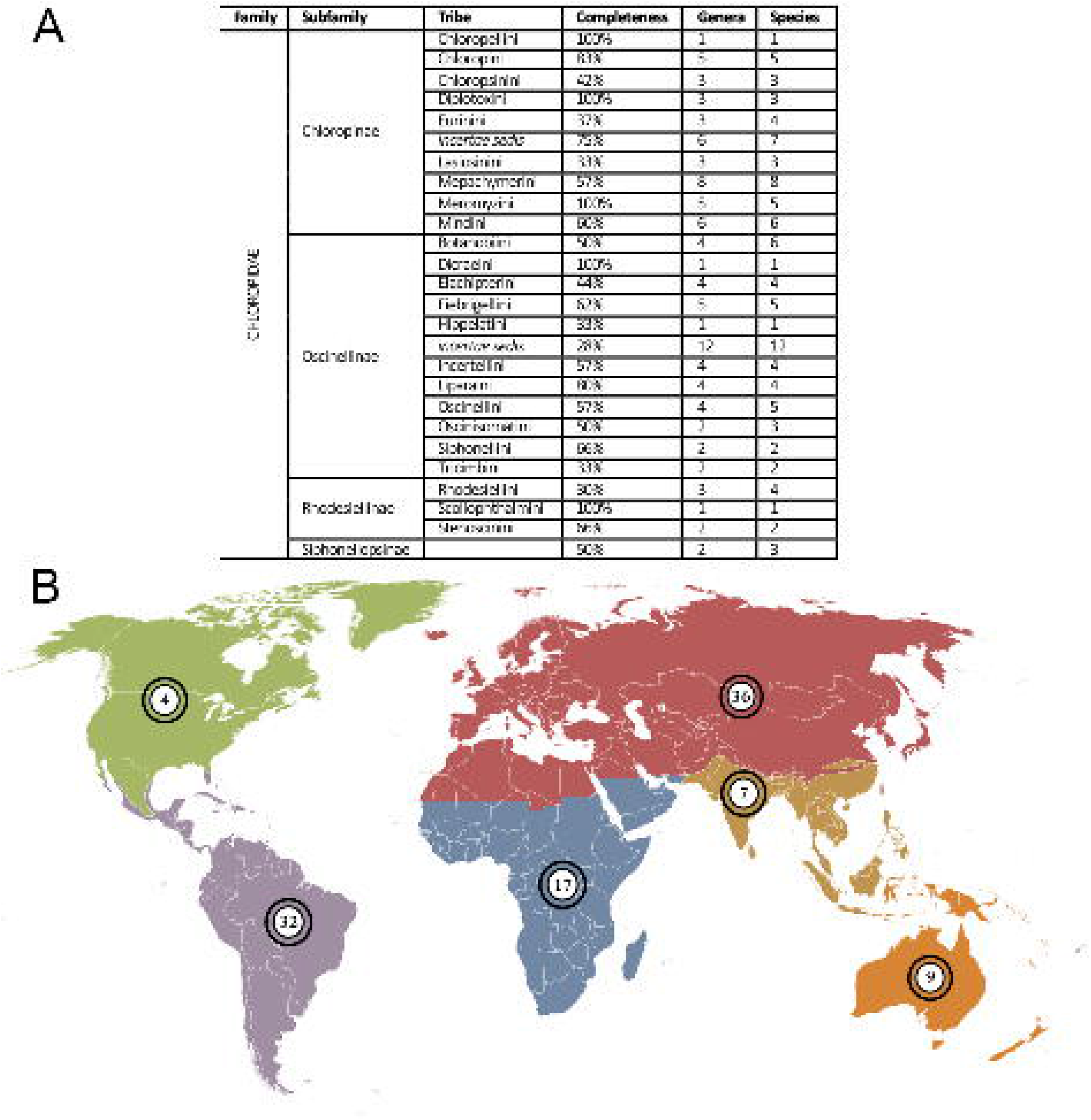
Traditional classification and distribution of the sampled species. The figure depicts the comprehensive taxon sampling and geographic coverage used in this study. **A**, Completeness, means the percentage of genera assigned to the tribe or more inclusive taxonomic rank that was used in this study; the two last columns represent the number of genera and species sampled per tribe respectively. **B**, Number of Chloropidae species per biogeographic region used in the phylogeny (image modified, By carol - Ecozones and Image:BlankMap-World6, compact.svg by User:Lokal_Profil., CC BY-SA 3.0, https://commons.wikimedia.org/w/index.php?curid=3698005). The nine species from Nepal were considered in the Palearctic fauna.

To test the phylogenetic affinities of Chloropidae within schizophoran flies, one family of Eremoneura and seven Schizophora families compose the outgroup—including Milichiidae and Neminidae previously recovered as sister to Chloropidae. Mitochondrial genes, UCEs and BUSCOs were newly generated for all taxa included in the datasets. Detailed information of taxon sampling is in Supplementary material 1.

The repositories acronyms are the following:

(ANHRT) African Natural History Research Trust, Kingsland, United Kingdom;

(ANIC) Australian National Insect Collection, Canberra, Australia;

(BMSA) National Museum Bloemfontein, Bloemfontein, South Africa;

(LEM) Lyman Entomological Museum, Sainte-Anne-de-Bellevue, Canada;

(MFN) Museum für Naturkunde Berlin, Berlin, Germany;

(MNHN) Museo Nacional de Historia Natural, Santiago, Chile;

(MNRJ) National Museum of Federal University of Rio de Janeiro, Rio de Janeiro, Brazil;

(MZUSP) Zoology Museum of Sao Paulo University, Sao Paulo, Brazil;

(NMC) National Museum Cardiff, Cardiff, United Kingdom;

(NMS) Stuttgart State Museum of Natural History, Stuttgart, Germany;

(SDEI) Senckenberg Deutsches Entomologisches Institut, Müncheberg, Germany;

(USNM) Smithsonian National Museum of Natural History, Washington DC, United States;

(ZSM) Zoologische Staatssammlung München, Munich, Germany.

### 2.2. Generation of genomic data

Non-destructive DNA extraction was performed using the DNAEasy kit (Qiagen) and digested overnight at 56°C following manufacturer protocol. The extracts were used as template for sequence the genome using whole-genome shotgun sequencing with low coverage. This method is suitable for retrieving data of both fresh samples and historical specimens that were included in our datasets. The DNA extracts were sent to Novogene GmbH (Munich, Germany) for genomic library preparation and sequencing. Sequencing was performed on an Illumina NovaSeq 6000 sequencer with 150bp paired-end reads.

### 2.3. Genomic data processing

The raw reads were initially deduplicated using Super Deduper (Peterson et al. 2015) before being trimmed and filtered using fastp (Chen et al. 2018). Overlapping paired-end reads were then merged using BBMerge to prepare them for de novo assembly (Bushnell et al. 2017). The genomes were then assembled de novo using Spades (Bankevich et al. 2012). The loci used for phylogenomic analyses were extracted following the methods outlined by Zhang (et al. 2019) and briefly discussed here.

Dataset 1 (Table S3) includes 15 protein-coding mitochondrial genes—ATP6, ATP8, COX1–3, CYTB, ND1–6, ND4L, rrnL, rrnS. The mitochondrial loci were extracted using MitoFinder (Allio et al. 2020). Both methods of extracting the mitochondrial genome using either directly using the filtered reads or the de novo assembly were attempted. The data resulting from the filtered reads were used as it was more likely to recover all mitochondrial genes in a single contig. We used a mitochondrial genome assembly from Chlorops oryzae (GenBank accession: MW438309) as a reference for MitoFinder. Alignment was conducted in MAFFT v7.475 (Katoh 2013) under E-INS-i strategy, except for the mitochondrial rRNA genes (Q-INS-i), and ND2 and ND6 (L-INS-i). Manual curation of the alignments was done in AliView (Larson 2014). Data matrix is available at Supplementary material 3.

UCEs were extracted from the de novo genome assemblies in silico using the pipeline outlined in phyluce. First, the probes for were designed based on the assembled genome of *Ceratobarys sacculicornis* (Enderlein), *Rhodesiella brimley* Sabrosky, *Apotropina rufithotax* (Duda), *Aphanotrigonum trilineatum* (Meigen), *Neohaplegis glabra* (Duda), and *Trigonomma* cf. *pubicollis* (Becker) which represent span the diversity of the tree (S2). Using the newly designed probes, we then extracted the associated UCE loci from the de novo assembled genome of each individual.

The BUSCO loci were extracted as part of the pipeline to assess genome assemblies (Simão et al. 2015). The BUSCO assessment was ran using the lineage dataset for diptera (odb10). Rather than simply producing quality assessment of each assembly, BUSCO uses AUGUSTUS to output a nucleotide sequence for each BUSCO locus (Stanke et al. 2006). Only complete, single-copy genes were retained for further analyses.

The complete genomic dataset contains a total of shared 7,690 loci comprising of 4,892 UCEs and 2,797 BUSCOs. To avoid duplicated markers, we removed 156 overlapping loci between BUSCOs and UCEs. Alignment was conducted in MAFFT v7.475 (Katoh 2013) under E-INS-i strategy; and trimming in TrimAl v1.4.1 (Capella-Gutiérrez et al 2009), parameters -automated1. We then selected the 50% gene occupancy matrix (117 taxa; 12.075 loci), and the 80% gene occupancy matrix (117 taxa; 1.618 loci), which were concatenated and analyzed using the script “CompileGenes.run” (Torres et al. 2022) in TNT v1.6 (Goloboff & Morales 2023). Additional phylogenetic summary statistics were calculated using AMAS v.0.94 (Borowiec 2016).

To reduce the computational burden and the effect of missing data, the 80% gene occupancy matrix (dataset 2) was chosen for downstream analyses. The dataset 2 is available at Supplementary material 4.

### 2.4. Phylogenetic analyses

We used ModelFinder (Kalyaanamoorthy et al. 2017) to select substitution models and partition scheme with the best fit. Maximum Likelihood (ML) analyses, including Ultrafast bootstrap (UFBoot) and SH approximate likelihood ratio (SH-support) test values, were performed in IQTree v.2 (Minh et al. 2020). Support was considered strong for UFBoot >= 99 and SH-support >= 98. The ML analyses were performed on datasets 1 and 2 separately and concatenated.

### 2.5. Data availability

Raw reads are available at at NCBI Sequence Read Archive (SRA) as a BioProject, accession number PRJNA1327883.

## 3. RESULTS

### 3.1. Dataset performance

We aimed to sequence 10X coverage for each individual and the resulting average coverage per sample was 8.06 ± 3.72X based on the reads mapped onto the assembly. The average assembly size is 237 ± 58Mb. This coverage allowed us to successfully retrieve the mitochondrial genome and UCEs (S1). On the other hand, the overall number of BUSCOs and their shared loci were significantly lower and variable when compared with UCEs (see Figure 2). The 80% gene occupancy matrix contains only nine BUSCO loci while the 50% matrix contains 2,712 BUSCOs. Better performance of UCEs over BUSCOs for low coverage sequencing was also demonstrated in Zhang et al. (2019). As expected, the individuals with less than 1,000 UCEs represent historical samples.

**Figure 2:**
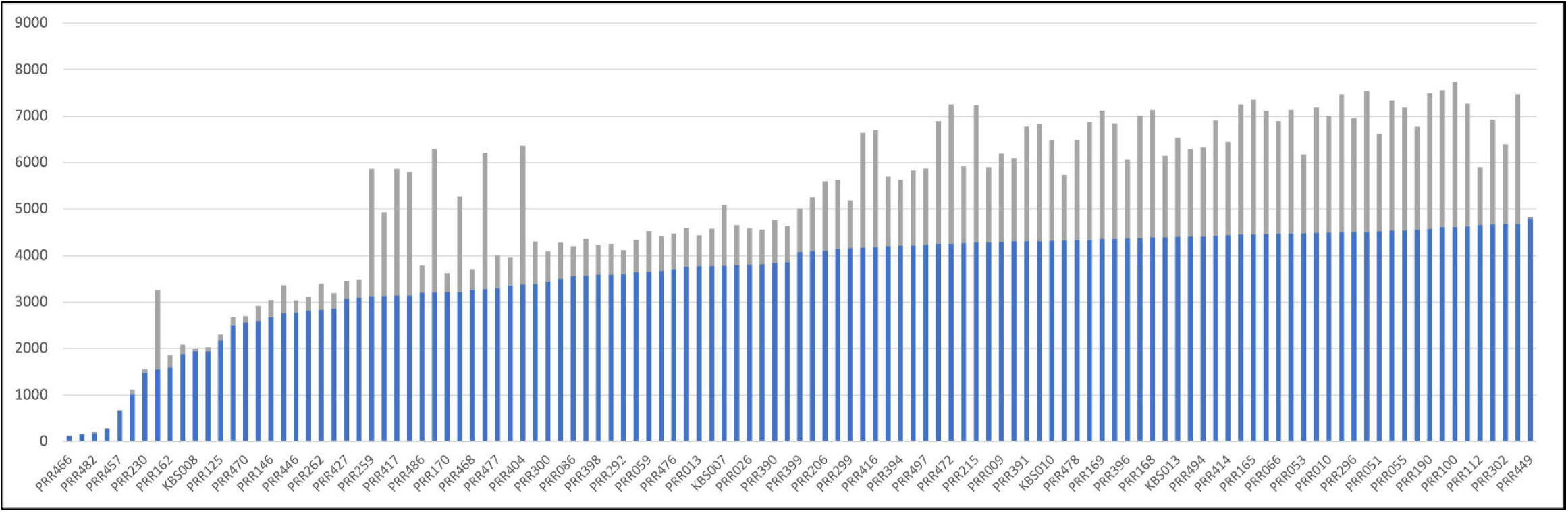
Comparison between number of UCEs and BUSCOs per sample in dataset 2 with 100% gene completeness. For all samples, UCEs are significantly more abundant than BUSCOs. Blue represents UCE loci and grey represent BUSCO loci.

Unlike the phylogenomic studies on flesh flies, dance flies, robber flies and shore flies and relatives (Buenaventura et al. 2020; Rhoden & Wahlberg 2020; Cohen et al. 2021; Winkler et al. 2022), our method skipped costs and lab workload associated with target enrichment by adopting whole-genome shotgun sequencing. Thallen (2022) used a similar approach to address Nereididae evolutionary relationships—another schizophoran lineage—with mitogenomes and 777 BUSCOs with 60.9% gene occupancy. Buenaventura (2020) used a specific bait set based on transcriptomes to retrieve protein-encoding UCEs. On the other hand, our UCE bait set was designed from genome assemblies, that enabled the capture of a broad diversity of nuclear loci.

Figure 4 presents a comparison between datasets 1 and 2. While the combination of UCEs and BUSCOs successfully resolved most deep nodes of the phylogeny, the mitogenome alone was insufficient to clarify the contentious position of Rhodesiellinae. Furthermore, the concatenated analysis of both datasets helped mitigate the rogue placement of two taxa with a high proportion of missing data. The ability to recover nearly 100% of the mitogenome across all species, even from historical samples, ensured an overlap of at least 14 loci among all taxa. Consequently, we selected the concatenated analyses (Figures 3–4) for interpreting the evolutionary relationships within Chloropidae and updating its classification system.

**Figure 3:**
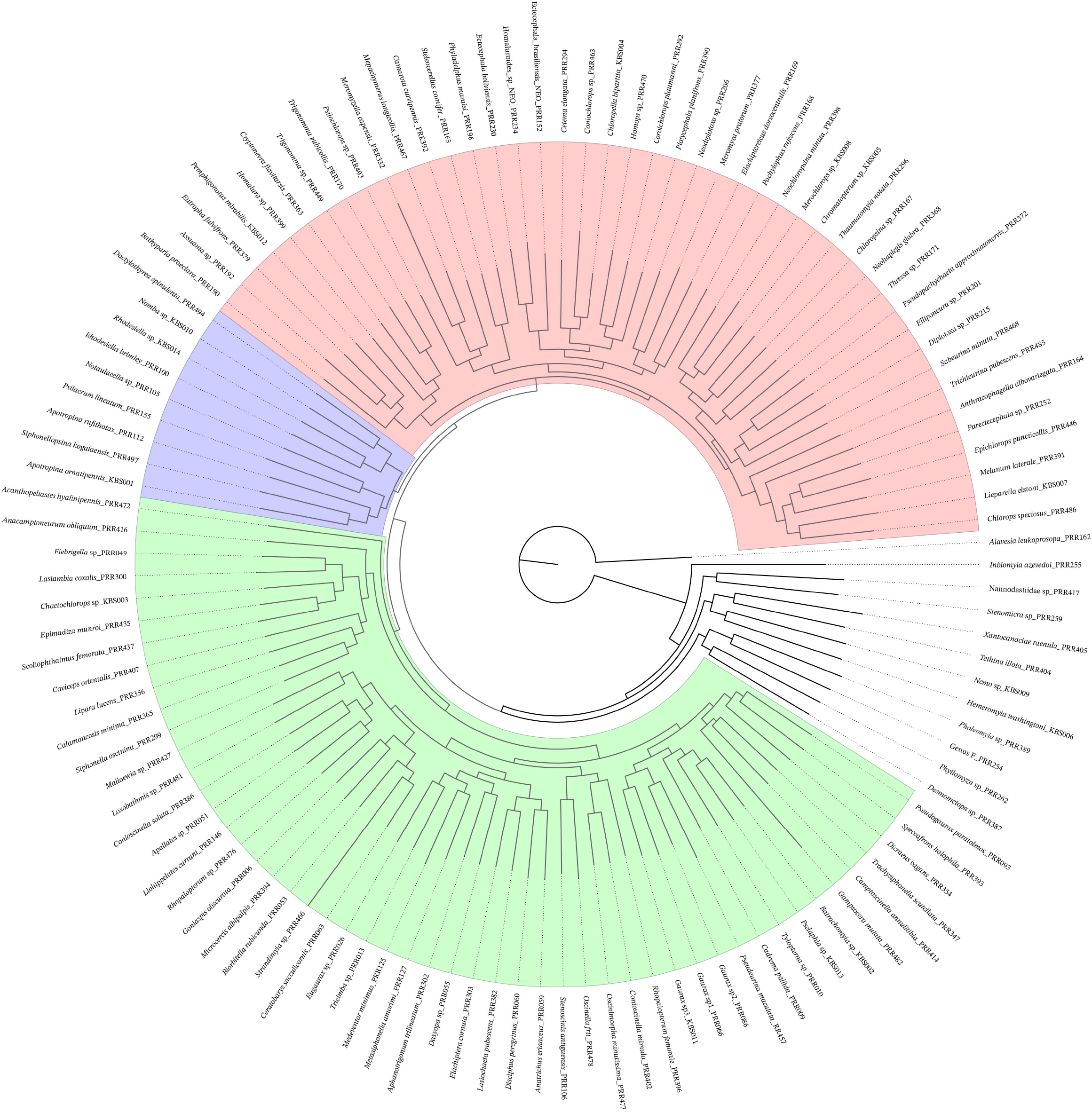
Maximum Likelihood tree of datasets 1 and 2 concatenated, showing the phylogenetic relationships among subfamilies of Chloropidae including the outgroup. Major clades are labeled according to subfamily-level classification proposed here. Red: Chloropinae; green: Oscinellinae and; purple: Siphonellopsinae.

**Figure 4:**
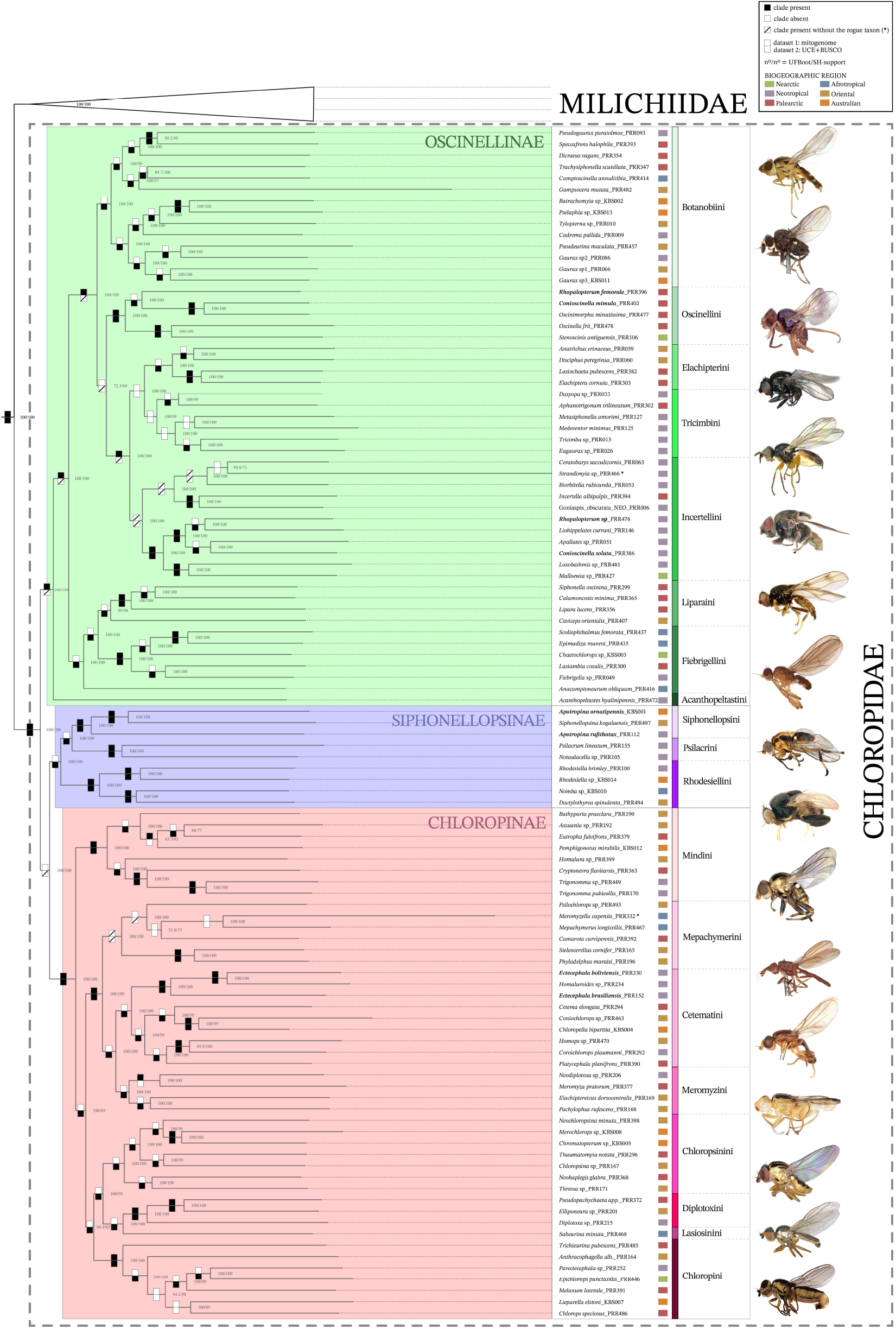
Maximum Likelihood (ML) tree of datasets 1 and 2 concatenated of Chloropidae+Milichiidae. The tree includes UFBoot and SH-Support values, thus the presence (black square)/absence (white square)/presence without rogue taxon (striped square) for each clade in the ML separate analyses. Clades are labeled according to subfamily-level and tribe-level classification proposed here. Red: Chloropinae; green: Oscinellinae and; purple: Siphonellopsinae. The biogeographic region of each sample follows Figure 1b color scheme; “*” = rogue taxon; taxon names “**in bold**” = genera recovered as non-monophyletic.

### 3.2. Delimitation of the Carnoidea and its position within Schizophora

Chloropidae has been regarded as the sister to the Milichiidae in most phylogenetic studies of cyclorrhaphan flies (e.g., McAlpine 1989; Buck 2006; Wiegmann et al. 2011), but not in Bayless et al. (2021), in which it appears as sister to the Neminidae based on transcriptomic data. Our results corroborate Chloropidae and Milichiidae as sister groups, and Neminidae as more closely related to Carnidae. It is worth noting that our taxon sampling encompasses all subfamilies of Milichiidae, including one undescribed genus mentioned by Swann (2010).

Although Song et al. (2022), using mitogenomic data, suggests that Chloropidae is embedded within Milichiidae, our results (Fig. 3) have 100% bootstrap and SH-support for the monophyly of both Chloropidae and Milichiidae as separate clades. Indeed, Bayless et al. (2021) and Song et al. (2022) include an extremely limited taxon sampling of these families, restricting their assessment of the relationships between Chloropidae and Milichiidae.

Buck’s (2006) delimitation of Carnoidea includes Acartophthalmidae, Australimyzidae, Canacidae, Carnidae, Chloropidae, Cryptochetidae, Inbiomyiidae, and Milichiidae. Our taxon sampling is not comprehensive enough for addressing the position of Carnoidea within Schizophora, but all Carnoidea families included—Canacidae, Carnidae, Chloropidae, and Milichiidae—except for Inbiomyiidae, were recovered in a clade with strong support (100% UFBoot and SH-support).

### 3.3. Morphological support for Chloropidae

As expected, Chloropidae was recovered as a monophyletic group (Figures 3–4). The oblique suture on the anepisternum of grass flies (Figure 5; arrow), first mentioned by Riccardi & Amorim (2020), revealed to be a synapomorphy of Chloropidae. A systematic investigation of this feature in schizophoran acalyptrate taxa, including the fossil taxon *Protoscinella electrica* Hennig, confirmed that the oblique suture is shared by all extant and extinct grass fly species, but it is absent in milichiids and other acalyptrates.

**Figure 5:**
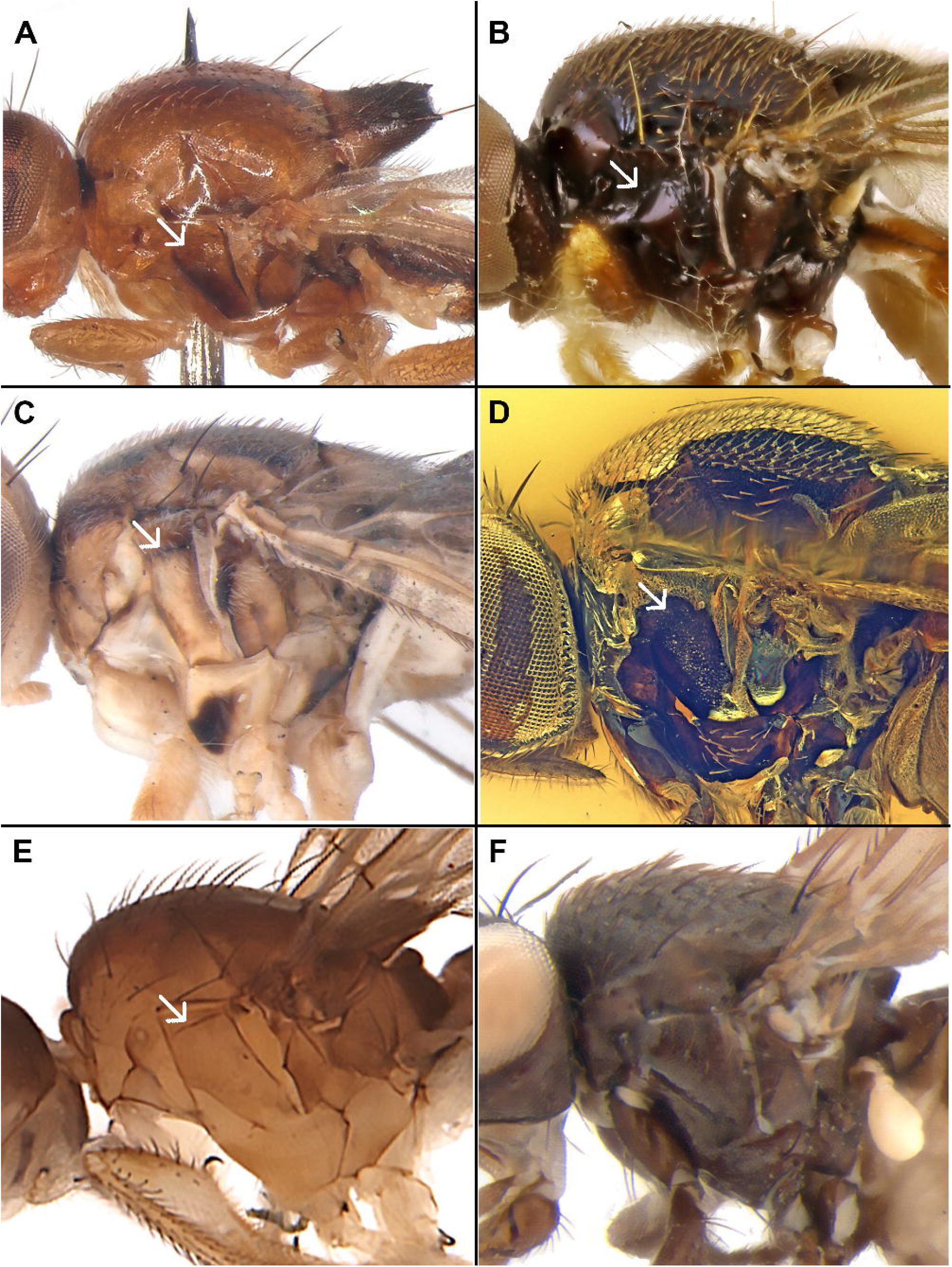
Thorax in lateral view. **A**, *Strandimyia* sp. (Chloropidae). **B**, *Acanthopeltastes curvinervis* (Chloropidae). **C**, *Tylopterna* sp. (Chloropidae). **D**, *Protoscinella electrica* (Chloropidae). **E**, *Apotropina rufithorax* (Chloropidae). **F**, *Meoneura flavifacies* (Carnidae). Arrow points to the oblique suture in the anepisternum.

MicroCT scans of *Protoscinella electrica* show that the oblique suture is actually a short apodeme in the pleuron. Judging by its position, it might be related to the direct flight muscles (see Dobi et al. 2015). The implications of this novel structure to the diversification and diversity of grass flies and its function, however, demands further investigation.

Unlike the propleural carina (Andersson 1977; Ismay et al. 2021), the oblique suture is easily seen in wet samples. The visualization of the oblique suture in dry grass flies with heavily pruinose pleuron is more difficult, although a depression on the anepisternum is always visible.

### 3.4. Classification of Chloropidae

Our results strongly support a reassessment of Chloropidae general classification system, including the relationships and boundaries at subfamily and tribe levels. The question of the Rhodesiellinae has been particularly more critical in the literature. The morphological phylogeny of the Oscinellinae of Bazyar (2019) failed to recover the rhodesielline genera *Psilacrum* and *Stenoscinis*as closely related to *Rhodesiella*.

We propose adjustments to the classification systems previously proposed by Nartshuk (2012), Mlynarek & Wheeler (2018), and Riccardi & Amorim (2020). A total of 19 genera, previously lacking tribal placement, are now incorporated into the grass fly classification system. While tribes in Chloropinae and Oscinellinae require minor adjustments, most Rodesiellinae genera are in fact nested with Siphonellopsinae. We propose here to expand the limits Siphonellopsinae to accommodate the rhodesielline genera, subdividing the subfamily into three tribes.

The inferred evolutionary relationships between grass fly subfamilies (Figures 3–4) are Oscinellinae+(Chloropinae+Siphonellopsinae). Overall, our study includes a new circumscription for the subfamily Siphonellopsinae, the proposal of three new tribes, new subfamily placement for seven genera, new tribe placement for 52 genera, revalidation of the tribe Cetematini, and the synonymy of Rhodesiellinae and three tribes.

The genera not sampled in this study are marked with * and remain here with the same classification status of previous systems or moved based on a well-established relationship with a sampled taxon.

#### Subfamily SIPHONELLOPSINAE Duda

Our phylogeny strongly supports the inclusion of taxa previously placed in Rhodesiellinae within Siphonellopsinae, rendering the former subfamily paraphyletic. Therefore, we propose Rhodesiellinae as a junior synonym of Siphonellopsinae **syn. nov**.

**Morphological diagnosis:** ocellar setae proclinate and costa extending to M1.

Further studies of the male genitalia and other morphological features are necessary to improve the morphological diagnosis of this clade.

Tribe Siphonellopsini Riccardi **trib. nov**.

**Type genus:** *Apotropina* Hendel

**Genera included:** *Apotropina* Hendel, *Siphonellopsina* Andersson, *Protohippelates* Andersson*, *Siphonellopsis* Strobl*.

**Comments**: The morphological diagnosis remains similar to previous diagnosis of Siphonellopsinae (see Andersson 1977) as 1–3 fronto-orbital setae lateroclinate, postpronotum with an inclinate seta; and asymmetrical pregenital segments in males. The type-species of *Apotropina*—*A. virduata* (Schiner)— is described from Australia (Nartshuk 2012). As *A. ornatipennis* (Malloch) from the Australian region is nested *Siphonellopsina kogalaensis* Andersson from the Oriental region, instead of with *A. rufithorax* (Duda) from the Neotropical region. Therefore, more sampling is necessary to resolve the boundaries and distribution of *Apotropina* within Siphonellopsini.

Tribe Psilacrini Riccardi **trib. nov**.

**Type genus:** *Psilacrum* Becker.

**Genera included:** *Psilacrum* Becker, *Notaulacella* Enderlein.

**Comments:** *Psilacrum* was previously assigned to the tribe Stenoscinini of Rhodesiellinae (Nartshuk 2012). However, our results and the diagnostic features—absence of anepisternal pilosity, along with the presence of a tibial organ and male cerci— described by Nartshuk (1983) as diagnostic for Stenoscinini are, in fact, widespread across grass flies.

The genus *Notaulacella* was previously placed in Oscinellinae without tribal assignment (Nartshuk 2012). However, all ML analyses indicate a close relationship between both genera.

#### Tribe Rhodesiellini Andersson

**Genera included:** *Rhodesiella* Adams, *Nomba* Walker, *Dactylothyrea* Meijere, *Amnonella* Cherian*, *Bharathella* Cherian*, *Euthyridium* Frey*, *Indophthalmus* Cherian*, *Neorhodesiella* Cherian*, *Pseudonomba* Cherian*, *Tachinoceros* Meijere*.

**Comments:** Unlike other grass fly genera with proclinate ocellars, the genera *Rhodesiella, Nomba, Dactylothyrea* and *Tachinoceros* (Andersson 1977) have ocellars highly divergent, almost forming a 180° angle, opposed to the usually divergent ocellars with less than 90°. This feature might be diagnostic, but it needs to be further explored in Rhodesiellini. The linear gena, wing with oblique dm-m and M_1_ with sinuosity posterior to dm-m are also diagnostic features of this tribe (Andersson 1977; Cherian 2002).

#### Subfamily CHLOROPINAE Rondani

Well-established lineage of grass flies (see Riccardi & Amorim 2020). This subfamily contains only changes in the tribal system. The tribe Chloropellini Riccardi & Amorim is nested within Cetematini, and genera previously assigned to the tribe Eurinini are scattered across the phylogeny.

#### Tribe Mindini Paramonov

**Genera included:** *Assuania* Becker, *Bathyparia* Lamb, *Cerais* Wulp*, *Collessimyia* Spencer*, *Cryptonevra* Lioy, *Eutropha* Loew, *Homalura* Meigen, *Neohaplegis* Beschovski*, *Pemphigonotus* Lamb, *Siphlus* Loew*, *Terusa* Kanmiya*, *Trigonomma* Enderlein.

**Comments:** Although *Eutropha* and *Bathyparia* were assigned to Mindini by Nartshuk (2012), their relationships were left as uncertain by Riccardi & Amorim (2020). Both genera were nested with *Trigonomma* in Riccardi & Amorim’s (2020) morphological phylogeny, the latter being without tribal placement until now.

#### Tribe Mepachymerini Nartshuk

**Genera included:** *Aragara* Walker*, *Bricelochlorops* Paganelli*, *Camarota* Meigen, *Centorisoma* Becker*, *Ischnochlorops* Paganelli*, *Mepachymerus* Speiser, *Metopostigma* Becker*, *Phyladelphus* Becker, *Psilochlorops* Duda, *Sagareocerus* Paganelli*, *Steleocerellus* Frey.

**Comments:** This is the first record of *Psilochlorops* from the Oriental region. The genus *Phyladelphus* was previously assigned to Lasiosinini (Riccardi & Amorim 2020; Nartshuk 2012). Despite *Meromyzella* is nested in Mepachymerini, we consider it as a rogue taxon due to the lack of support and unstable position even in preliminary analyses.

#### Tribe Cetematini Nartshuk

**Genera included:** *Bothynocerus* Paganelli*, *Cetema* Hendel, *Chloropella* Malloch, *Coniochlorops* Duda, *Coroichlorops* Paganelli, *Ectecephala* Macquart, *Homaluroides* Sabrosky, *Homops* Speiser, *Platycephala* Fallen.

**Comments:** The tribe Cetematini was originally established to accommodate the genera *Cetema* and *Homaluroides* (Nartshuk 1983), both characterized by a large ocellar triangle and phytophagous larvae. Notably, *Ectecephala* and *Platycephala* also possess phytophagous larvae and, along with *Homaluroides*, exhibit obligate viviparity (Ismay et al. 2021; Riccardi 2012; Meier et al. 1999). Our phylogeny supports the inclusion of the genera previously placed in the Chloropellini within the Cetematini, a clade in this analysis with 100% support. We formally propose here Chloropellini as a junior synonym of Cetematini **syn. nov**. The boundaries between *Ectecephala, Ectecephalina* Paganelli and *Homaluroides* are still obscure (Riccardi & Amorim 2020; Riccardi 2012). *Ectecephalina* has been previously considered a junior synonym of *Ectecephala* (Riccardi 2012). The monotypic *Bothynocerus* form well-established clade (Riccardi & Amorim 2020). Therefore, *Bothynocerus* is moved from Eurinini to this tribe.

#### Tribe Meromyzini Nartshuk

**Genera included:** *Chloromerus* Becker*, *Elachiptereicus* Becker, *Meromyza* Meigen, *Neodiplotoxa* Malloch, *Pachylophus* Loew.

**Comments:** *Elachiptereicus* was previously regarded as Mepachymerini (Riccardi & Amorim 2020) and fits well into the Meromyzini. *Neodiplotoxa* is confirmed as sister to *Meromyza*. The male genitalia of Meromyzini genera present multiple modifications. Pre- and postgonites are fused in *Neodiplotoxa*, phallic complex extremely elongated in *Elachiptereicus*, and postgonites orientation and complexity are increased in *Meromyza*. The male genitalia of *Meromyzella* is similar to *Chloromerus* and *Pachylophus* (see Riccardi & Amorim 2020). Fresh samples of *Meromyzella* would clarify the position of this rogue taxon.

#### Tribe Chloropsinini Riccardi & Amorim

**Genera included:** *Chloropsina* Becker, *Chromatopterum* Becker, *Cordylosomides* Strand*, *Ensiferella* Andersson*, *Merochlorops* Howlet, *Neochloropsina* Riccardi & Amorim, *Neoloxotaenia* Sabrosky*, *Neohaplegis* Beschovski, *Pseudochromatopterum* Deeming*, *Thressa* Walker, *Thaumatomyia* Zenker.

**Comments:** *Thaumatomyia* was one of the rogue taxa in Riccardi & Amorim (2020), and not assigned to any tribe. *Merochlorops* is transferred from Mindini to the Chloropsini.

#### Tribe Diplotoxini Nartshuk

**Genera included:** *Diplotoxa* Loew, *Elliponeura* Loew, *Pseudopachychaeta* Strobl.

**Comments:** Similar to the clade recovered by Riccardi & Amorim (2020).

#### Tribe Lasiosinini Nartshuk

**Genera included:** *Desertochlorops* Nartshuk*, *Lagaroceras* Becker*, *Lasiosina* Becker*, *Sabeurina* Deeming, *Stenophthalmus* Becker*, *Urubambina* Paganelli*.

**Comments:** After the transfer of *Phyladelphus* to Mepachymerini and *Assuania* to Mindini, the only sampled genus from this tribe is *Sabeurina*. Additional sampling is needed to verify the status of Lasiosinini, what will allow assessing whether the complete fusion of the surstylus to the epandrium represents a unique event in Chloropinae evolution.

#### Tribe Chloropini Rondani

**Genera included:** *Anthracophagella* Andersson, *Capnoptera* Loew*, *Chlorops* Meigen, *Dudeurina* Ismay*, *Epichlorops* Becker, *Eurina* Meigen*, *Lieparella* Spencer, *Melanum* Becker, *Parectecephala* Becker.

**Comments:** The unsampled genera *Eurina* and *Dudeurina* are included in this tribe due to their close affinities with *Trichieurina* (see Riccardi & Amorim 2020). The monophyly of the worldwide genus *Chlorops* has been questioned before (Riccardi & Amorim 2020; Ismay et al. 2021). A large sampling of species of this genus is still necessary to clarify the boundaries of the genus and how other Chloropini genera are nested within *Chlorops*. This will also allow properly recovering the phylogenetic signal of some morphological features within the tribe— pre- and postgonites parallel, palpus with longitudinal sulcus, postpedicel length, pruinosity and coloration patterns, ocellar triangle striation, and arista pilosity.

#### Subfamily OSCINELLINAE Becker

It will still be necessary, along the next years, to expand the oscinelline taxonomic sampling to have a wider phylogenetic coverage of the subfamily with molecular data. This analysis shows that *Stenoscinis* Malloch and *Scoliophthalmus* Becker, previously assigned to Rhodesiellinae (Nartshuk 2012), clearly fits in the Oscinellinae (see Figure 4). The proclinate ocellar setae present in these genera, hence, should be seen as homoplastic since most oscinellines have erect or reclinate ocellars. Moreover, the tribes Dicraeini Nartshuk and Siphonellini Lioy are not supported by our analyses.

Tribe Acanthopeltastini Riccardi **trib. nov**.

**Type genus:** *Acanthopeltastes* Enderlein.

**Genera included:** *Acanthopeltastes* Enderlein, *Agrophaspidium* Wheeler & Mlynarek*.

**Comments:** The genus *Acanthopeltastes* is endemic to South America and has a distinctive morphology, including long, fingerlike projections on the scutellum and a unique head shape characterized by kidney-shaped eyes and a high ventral facial margin. On the other hand, *Agrophaspidium* is restricted to Central America, and also exhibits scutellar extensions and a distinctive male genitalia (Wheeler & Mlynarek 2008). The phylogenetic placement of *Agrophaspidium* within the Oscinellinae has been uncertain, while morphological features suggest it may be close to *Acanthopeltastes*.

#### Tribe Fiebrigellini Nartshuk

**Genera included:** *Anacamptoneurum* Becker, *Chaetochlorops* Malloch, *Epimadiza* Becker, *Fiebrigella* Duda, *Heteroscinis* Lamb*, *Heteroscinoides* Cherian*, *Lasiambia* Anonymous, *Polyodaspis* Duda*, *Scoliophthalmus*.

**Comments:** The placement of *Scoliophthalmus* in this tribe as sister to *Epimadiza* has strong support.

#### Tribe Liparaini Camero & Tubbs

**Genera included:** *Anomoeoceros* Lamb*, *Calamoncosis* Enderlein, *Caviceps* Malloch, *Lipara* Meigen, *Siphonella* Macquart.

**Comments:** Our phylogeny support the inclusion of taxa previously placed in Siphonellini within Liparaini. Therefore, we propose Siphonellini as a junior synonym of Liparaini **syn. nov**. Some morphological features as shiny and robust body are shared by the clade Fiebrigellini+Liparaini.

#### Tribe Incertellini Nartshuk

**Genera included:** *Apallates* Sabrosky, *Biorbitella* Sabrosky, *Ceratobarys* Coquillett, *Conioscinella* Duda, *Goniaspis* Duda, *Incertella* Sabrosky, *Liohippelates* Duda, *Loxobathmis* Enderlein, *Malloewia* Sabrosky, *Meijerella* Sabrosky*, *Microcercis* Beschovski*, *Parameijerella* Cherian*, *Parapallates* Cherian & Tilak*, *Rhopalopterum* Duda, *Strandimyia* Duda.

**Comments:** The type species of *Rhopalopterum—R. limitatum* (Becker)*—*is from Haiti and the Neotropical species assigned to this genus are considered congeneric. A taxonomic revision of the genus, nevertheless, is needed to properly set its taxonomic boundaries. The Neotropical *Conioscinella soluta* (Becker) (type species of *Conioscinella*), included in our analysis, is shown to belong in the Incertellini and the genus is transfered from the Oscinellini. *Conioscinella* as well requires a taxonomic revision. *Ceratobarys* and *Goniaspis* are transferred from Elachipterini, while *Strandimyia* and *Loxobathmis*, which had no previous tribal assignment in the Oscinellinae, are shown to fit into the Incertellini. The relationships between *Ceratobarys* and *Strandimyia* should be treated carefully as the latter behaved as rogue in our analyses.

#### Tribe Tricimbini Nartshuk

**Genera included:** *Aphanotrigonella* Nartshuk*, *Aphanotrigonum* Duda, *Aprometopis* Becker*, *Dasyopa* Malloch, *Eugaurax* Malloch, *Indometopis* Cherian*, *Metasiphonella* Duda, *Medeventor* Wheeler, *Pseudotricimba* Ismay*, *Siphunculina* Rondani*, *Tricimba* Lioy, *Tricimbomyia* Cherian*.

**Comments:** This tribe previously included only *Tricimba. Dasyopa* is transferred from Liparaini, and *Aphanotrigonum* from Incertellini. Moreover, *Eugaurax, Metasiphonella* and *Medeventor* had no previous tribal status.

#### Tribe Elachipterini Lioy

**Genera included:** *Allomedeia* Mlynarek & Wheeler*, *Alombus* Becker*, *Anatrichus* Loew, *Disciphus* Becker, *Elachiptera* Macquart, *Lasiochaeta* Corti, *Melanochaetomyia* Cherian*, *Sepsidoscinis* Hendel*.

**Comments:** The position of *Ceratobarys* away from the Elachipterini is quite surprising. This result reinforces how plastic morphological features as enlarged arista and elongated body might be to (Riccardi & Amorim 2016; Andersson 1979). As no species previously assigned to *Cyrtomomyia* Becker and *Togeciphus* Nishijima were included in our sampling. We hence follow Mlynarek & Wheeler’s (2018) position, considering both genera as junior synonyms of *Elachiptera*. In contrast, the previous placement of *Ceratobarys* and *Goniaspis* within Elachipterini (Mlynarek & Wheeler 2018) is not supported by our results. The controversial tribal assignment of these genera may be related to limited taxon sampling within Oscinellinae (Mlynarek & Wheeler 2018) and difficulties in interpreting plastic morphological characters, such as thickened arista and reduced anal lobe of the wing. Moreover, *Goniaspis* was treated as a rogue taxon by Bazyar (2019), which indicates uncertainty in the phylogenetic relationships of this genus within Oscinellinae based on morphology.

#### Tribe Oscinellini Becker

**Genera included:** *Neolcella* Cherian*, *Neoscinella* Sabrosky*, *Olcella* Enderlein*, *Oscinella* Becker, *Oscinimorpha* Lioy, *Stenoscinis* Malloch.

**Comments:** The species *Conioscinella mimula* is nested with *Oscinimorpha* which indicates further taxonomic revisions required for these genera in the Palearctic region. Bazyar (2019) results also recovered *Stenoscinis* nested with other Oscinellinae genera, although with a different clade composition. The only congruence with our results is the close relationship between *Rhopalopterum* and *Stenoscinis*, which reinforces the need to revise the taxonomic assignment of the 20 valid species of *Rhopalopterum* distributed in the Nearctic, Neotropical and Palearctic regions (von Tschirnhaus & Groll 2024).

#### Tribe Botanobiini Malloch

**Genera included:** *Batrachomyia* Krefft, *Cadrema* Walker, *Camptoscinella* Sabrosky, *Cestoplectus* Lamb*, *Dicraeus* Loew, *Gaurax* Loew, *Gampsocera* Schiner, *Hapleginella* Duda*, *Leucochaeta* Becker*, *Pselaphia* Becker, *Pseudeurina* Meijere, *Pseudogaurax* Malloch, *Pterogaurax* Duda*, *Trachysiphonella* Enderlein, *Tylopterna* Bezzi.

**Comments:** Our phylogeny support the inclusion of taxa previously placed in Dicraeinni within Botanobiini. Therefore, we propose Dicraeinni as a junior synonym of Botanobiini **syn. nov**. The genera *Batrachomyia, Cadrema, Trachysiphonella* and *Tylopterna* have no previous tribal status and *Pseudeurina* is transferred from Liparaini.

#### Tribe Hippelatini Duda

**Genera included:** *Chaethippus* Duda*, *Hippelates* Loew*, *Lioscinella* Duda*.

**Comments:** With the transfer of *Liohippelates* to Incertellini, this tribe is not sampled.

#### Tribe Oscinisomatini Enderlein

**Genera included:** *Eribolus* Becker*, *Oscinisoma* Lioy*, *Sabroskyina* Beschovski*.

**Comments:** With the transfer of *Rhopalopterum* to Incertellini, this tribe is not sampled.

*Incertae sedis*—without tribal assignment

**Genera included:** *Arcuator* Sabrosky*, *Aulacogaurax* Meijere in Becker & Meijere*, *Benjaminella* Malloch*, *Camptopeltes* Mlynarek & Wheeler*, *Cauloscinis* Yang & Yang*, *Coryphisoptron* Enderlein*, *Deltastoma* Malloch*, *Discograstella* Enderlein*, *Dysartia* Sabrosky*, *Enderleiniella* Becker*, *Indonella* Cherian in Cherian & Shinimol*, *Kurumemyia* Kanmiya*, *Kwarea* Sabrosky*, *Merobates* Duda*, *Merodonta* Malloch*, *Mimosepsis* Sabrosky*, *Monochaetoscinella* Duda*, *Onychaspidium* Enderlein*, *Opetiophora* Loew*, *Oscinicita* Wheeler*, *Paraapallates* Cherian in Cherian & Tilak*, *Parasiphonella* Enderlein*, *Platyina* Malloch*, *Pseudogampsocera* Sabrosky*, *Sacatonia* Sabrosky*, *Siphonellomyia* Seguy*, *Thyridula* Becker*, *Tropidoscinis* Enderlein*, *Vanchium* Cherian*.

## 4. FINAL REMARKS

This study generated genomic resources from all chloropid subfamilies, 95% of the former chloropid tribes and 48% of the chloropid genera across all biogeographic regions. This sampling is considerably comprehensive and allows us to critically evaluate existing classification systems and resolve some of the long-standing obscure relationships within the Chloropidae. This, as quite expected, led to quite dramatic implications for the grass fly classification system. Based on the phylogeny, we redefine Siphonellopsinae, propose two new Siphonellopsinae and one Oscinellinae tribes, revalidate one tribe, reassign the tribal position of 59 genera, and synonymize four suprageneric taxa.

The resolution of deep nodes here largely supported by UCE data. This yielded the first Chloropidae backbone phylogeny, which is key to interpreting the evolution of morphological and biological traits in the family as their plasticity. In fact, it provides a baseline for grass fly biodiversity and evolution studies worldwide, such as understanding the ecological diversity (Riccardi & Hartop 2024; Matallana-Puerto et al. 2025) and evolutionary history of grass fly unique traits (Riccardi et al. 2024).

The sampling of the two most speciose subfamilies—Chloropinae and Oscinellinae—, despite the considerably wide taxonomic coverage, is limited to confidently resolve additional taxonomic conundrums: 29 Oscinellinae genera are still left as *incertae sedis*, while the comprehensive overview of Chloropinae provided by Riccardi & Amorim (2020) has been instrumental to assign tribal status to virtually all chloropine genera.

Species of *Apotropina, Conioscinella, Ectecephala* and *Rhopalopterum* are rather scattered and makes evident the need of taxonomic revisions for these genera across their entire geographic ranges. Notably, *Conioscinella* and *Ectecephala* have been suggested as potentially paraphyletic (Riccardi & Amorim 2020; Ismay et al. 2021). Additionally, the two Oriental and Australian Siphonellopsini genera appear more closely related to each other than to their corresponding Neotropical species, what stresses the urgent need of a comprehensive investigation of *Apotropina*.

The phylogenetic relationships recovered here set a starting point to support comprehensive taxonomic studies of chloropid genera with known problematic boundaries (Riccardi & Amorim 2020; Riccardi & Hartop 2024; Foster 2024). Resolving their contentious taxonomy is fundamental to accelerating the species discovery process and, consequently, to overcoming the taxonomic neglect of this hyperdiverse lineage.

*Meromyzella* and *Stradimyia* are represented by historical specimens. These taxa exhibit long branches in the phylogeny, likely due to a high proportion of missing data (S1). They behave as rogue taxa (Figure 4) and their position should still be interpreted carefully. In dataset 2 analysis, *Stradimyia* formed a polytomy with Siphonellopsinae and Chloropinae, whereas the mitogenome and concatenated analyses placed it with *Ceratobarys* in Oscinellinae. Morphological traits, including erect and cruciate ocellar setae and the structure of the surstylus, support its placement in Oscinellinae.

The position of *Meromyzella* was more consistent in all analyses, nested within Chloropinae. It was recovered as sister to *Pachylophus* (dataset 1) and to *Meromyza* (dataset 2), both in Meromyzini, but appeared as sister to *Mepachymerus* (Mepachymerini) in the concatenated analysis. The male genitalia of *Meromyzella* resemble those of Meromyzini taxa (see Riccardi & Amorim 2020). Both *Meromyzella* and *Stradimyia* are monotypic and rare in collections. *Stradimyia* is restricted to the Neotropical region, while *Meromyzella* is found in the Afrotropical region (von Tschirnhaus & Groll 2024). Data from fresh specimens, along with a systematic study of male genitalia across Chloropinae and Oscinellinae, should resolve the ambiguous position of these two genera.

Carnoidea and Opomyzoidea are poorly defined superfamilies of Schizophora (McAlpine 1989; Buck 2006). Although the relationships close to the root in phylogenetic analyses have lower reliability, the controversial placement of Inbiomyiidae and Neminidae in our topology (Figure 3), suggest that a more comprehensive taxon sampling of Carnoidea families, particularly of Acartophthalmidae, Australimyzidae and other Opomyzoidea families, would be necessary to clarify the position, circumscription and relationships between these superfamilies within the Schizophora.

Similar to our results for dataset 1, Liu et al. (2024) also recovered Chloropidae as monophyletic using mitogenomes, but they were not able to confidently recover deep nodes in grass fly phylogeny, possibly due to the rapid diversification of Schizophora. Although with a limited sampling, Li et al. (2017) recovered traditional relationships between families of Lauxanioidea using mitogenomes, but failed to reveal higher-level relationships between schizophoran superfamilies. On the other hand, mitogenomes could provide new insights into evolutionary history of Chironomidae (Lin et al. 2022). The inclusion of nuclear loci (dataset 2) mainly based on UCEs was fundamental for resolving deep nodes in grass fly phylogeny, also corroborating the utility of UCEs in Diptera phylogenomics (Zhang et al. 2019; Buenaventura 2020) and limitations of mitogenomes alone to yield phylogenetic signal to fully recover the phylogenetic relationships within the Schizophora backbone.

Solving the problem of the uncertain position of Rhodesiellinae, setting the relationships between all chloropid subfamilies, and validating a tribal system for the entire family could be successfully achieved in this study based on a phylogeny with robust support. It is clear that the concatenated analysis here including UCEs, BUSCOs and the mitogenome provide better results than separate analyses (Figure 4), as it supports the chloropid backbone while mitigates the effect of missing data.

Overall, our methodological approach was successful for retrieving thousands of UCEs and the mitogenome while preserving vouchers for morphological study and skipping costs associated with target enrichment. Moreover, the bait set generated (S2) was effective for extracting UCEs from both Schizophora families (Canacidae, Carnidae, Chloropidae, Inbiomyiidae, Milichiidae, Nannodastiidae, Neminidae and Periscellidae) and for Eremoneura (Atelestidae). These outputs indicate that our strategy could provide a novel framework for understanding the evolution of the last and most species-rich radiation of flies.

## ACKNOWLEDGEMENTS

PRR benefitted from the CAPES-HUMBOLDT fellowship (88881.512934/2020-01). This publication benefited from the HPC Service of FUB-IT, Freie Universität Berlin, for computing time (10.17169/refubium-26754). We are grateful to Christel Hoffeins for lending inclusions of *Protoscinella electrica*, to John and Barbara Ismay for donating specimens, and to Jessica Gillung (LEM), Jung Kim (USNM), Frank Menzel (SDEI), John Deeming (NMC), Ashley Kirk-Spriggs (ANHRT), Burgert Muller (BMSA), and Daniel Whitmore (NMS) for lending specimens to conduct this project.

## AUTHOR CONTRIBUTION

**Paula Raile Riccardi:** conceptualization, data curation, visualization, funding acquisition, formal analysis (lead), methodology (equal), project administration (lead), resources (lead), writing – original draft/review & editing (lead). **Dalton de Souza Amorim:** resources (supporting), review & editing (supporting). **Keith Bayless:** resources (supporting). **Joshua V. Peñalba:** formal analysis (supporting), methodology (equal), project administration (supporting), writing – original draft/review & editing (supporting).

